# Teaching good agronomic practices to strengthen pigeon pea cultivation and value chains through farmer-to-farmer education in Zambia

**DOI:** 10.1101/2025.06.07.657896

**Authors:** Hamid Khazaei, Sebastian Scott, Jarkko K. Niemi

## Abstract

Farmer field schools (FFSs) are transformative, participatory approaches to adult agricultural education that effectively promote farmers’ learning and capacity building and empower farming communities with practical knowledge. Here, we present a case study of FFS in the Katete district, Eastern province of Zambia, focused on good pigeon pea agronomic practices and the value chain. Three FFSs were established, focusing on a variety of trials, bio-pesticides, and a pigeon pea-specific rhizobial inoculant. Extreme weather conditions, an underdeveloped pigeon pea seed value chain, and a poor seed system hindered the implementation of the FFSs. To strengthen the pigeon pea seed value chain, a seed dehuller is now available to farmers. Our results highlight the importance of the design, implementation, monitoring, and evaluation of the FFSs. Pigeon pea FFSs will support crop and food diversification, improve soil fertility and sustainable agriculture, and may increase household income.

## INTRODUCTION

Farmer field schools (FFSs) are initiatives where smallholder farmers establish and manage demonstration plots to showcase agricultural practices and technologies aimed at enhancing resilience, productivity, profitability, and household nutrition security. Given the positive impact FFS can have on rural livelihoods, they hold substantial potential to contribute to the achievement of the United Nations sustainable development goals (van den Berg et al., 2020). The farmers’ schools are established and managed in a way that enables them to clearly observe the tangible benefits of implementing good agronomic practices within their cropping systems. The FFS is a participatory, peer-to-peer learning approach designed to educate farmers in an informal setting within their own environment (FAO, 2001). The FFS is a widely adopted education and extension approach that employs experiential learning and group-based methods that support farmers in decision-making, problem-solving, and adoption of new techniques (Davis et al., 2012).

The FFS has been implemented and adapted globally, including Africa (e.g., Asiabaka, 2002; Duveskog et al., 2011; Davis et al., 2012; van den Berg et al., 2021a and 2021b; Pienaah et al., 2024). In Zambia, FFSs were first established in the mid-1990s under the Farm-Level Applied Research Methods Programme for Eastern and Southern Africa (FARMESA) initiative, which was introduced by the FAO to support Eastern and Southern Africa (Anandajayasekeram et al., 2007). A few more projects introduced FFS to Zambia in the early 2000s, all of which were supported by the FAO (Braun et al., 2006). Building on these early efforts, a decade later an FFS was established in Zambia’s Eastern Province with support from the World Bank’s BioCarbon fund initiative for sustainable forest landscapes (World Bank, 2019). In recent years, several other FFS programs affiliated with both the government and NGO sectors have also been developed, with varying focus areas and methodologies. These activities include Strengthening the Climate Resilience of Agricultural Livelihoods in Agroecological Regions I and II in Zambia (SCLARA, Ministry of Agriculture Zambia, 2018), SelfHelp Africa (2014), and World Vision Zambia (World Vision, 2021). FFS are well-suited to increase the capacity of smallholders to successfully cultivate, utilise and market new crops such as pigeon pea. Introduction of a new crop to farmers supports the adoption of crop and diet diversification strategies by the Zambian government to reduce the risks linked to heavy reliance on crop monocultures (Mangaba, 2017).

Pigeon pea (*Cajanus cajan* [L.] Millspaugh) is a promising grain legume drought adapted crop for the Eastern province of Zambia, with efforts underway to promote its cultivation. It has great potential for use as food, income generation, livestock feed and agroforestry as a soil fertility-improving crop. Pigeon pea is traditionally consumed in small quantities in Zambia as a fresh vegetable or as a whole dry grain. Cooking whole dry grain requires substantial cooking time and ranks poorly against alternative legumes, such as cowpea and common beans. Global annual pigeon pea production was around 4.6 million tonnes (Tg), cultivated across nearly 5.3 Mha in 2023. India was the largest producer, contributing 76% of the world’s production, followed by Myanmar (8%), Kenya (6%), Malawi (5%), and the United Republic of Tanzania (4%) (FAOSTAT, 2025). In India, the major producer and consumer of pigeon pea, its dry, split, and dehulled seeds, are called ‘dahl’ and are used as food (Locali-Pereira et al., 2023). In the semi-arid tropics, particularly during the dry season, the deep-rooted pigeon pea crop serves as insurance against drought (Sameer Kumar et al., 2017) while nurturing the soil for other crops as a great symbiotic nitrogen-fixer grain legume crop (Peoples et al., 2021). Its seeds are rich in protein (21–28% seed dry weight) along with vital amino acids and vitamins (Gomezulu and Mongi, 2022). Pigeon pea is a climate-smart crop that combines optimal nutritional profiles, high adaptation to environmental stresses, high biomass productivity, and moisture contributions to the soil. The increasing Zambian local and global export demands for this versatile legume present an opportunity to increase productivity, enhance seed delivery systems, and initiate value chains. These factors are making the crop increasingly important in the Eastern province of Zambia.

Although FFSs could help to valorize pigeon pea among smallholders, there has been no pigeon pea-based FFS, particularly focused on good agronomic management practices and value chain development in Zambia. The key issues for pigeon pea production in the region include being a minor crop (Phiri et al., 2024), a lack of a value chain, a limited number of adapted high-yielding germplasm, pest management, free-ranging livestock, and poor seed systems. Thus, FFSs were established in the Katete region of Zambia’s Eastern Province aims to increase awareness of pigeon pea crops, highlight their benefits, promote their cultivation via good agronomic practices, and support the development of a value chain by introducing dehulling equipment for seed processing to farmers. Here, we share lessons learned from these efforts, including the challenges encountered, opportunities identified, and strategies implemented to strengthen pigeon pea production and utilization.

## METHODOLOGY

The FFS for improved pigeon pea agronomy was carried out with three communities (Chimutende, Makwenda, and Greya; **Figure 1**) in the Katete region that already had an established FFS program running with Sebastian Scott of Grassroots Trust Limited in Zambia. The project focused on the pigeon pea value chain and gave rise to three new FFSs, with the goal of educating farmers on the main constraints for pigeon pea and possible solutions. **Table 1** outlines key challenges in pigeon pea cultivation in Zambia and presents targeted opportunities and responses implemented through FFSs.

**Figure 1.**
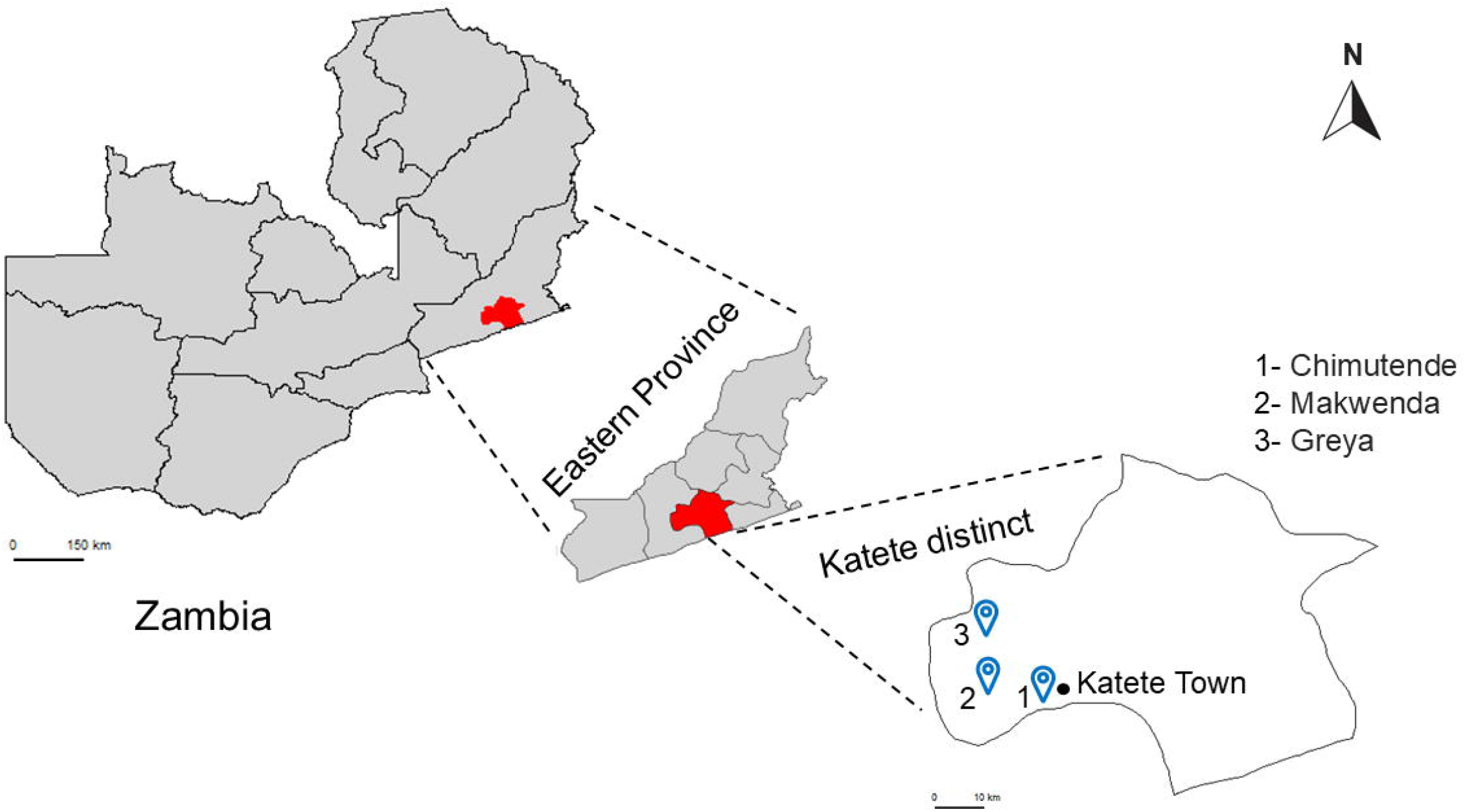
Map of Zambia showing the study area of three FFSs in Katete district, Eastern province. Map source: https://gadm.org/.

**Table 1.**
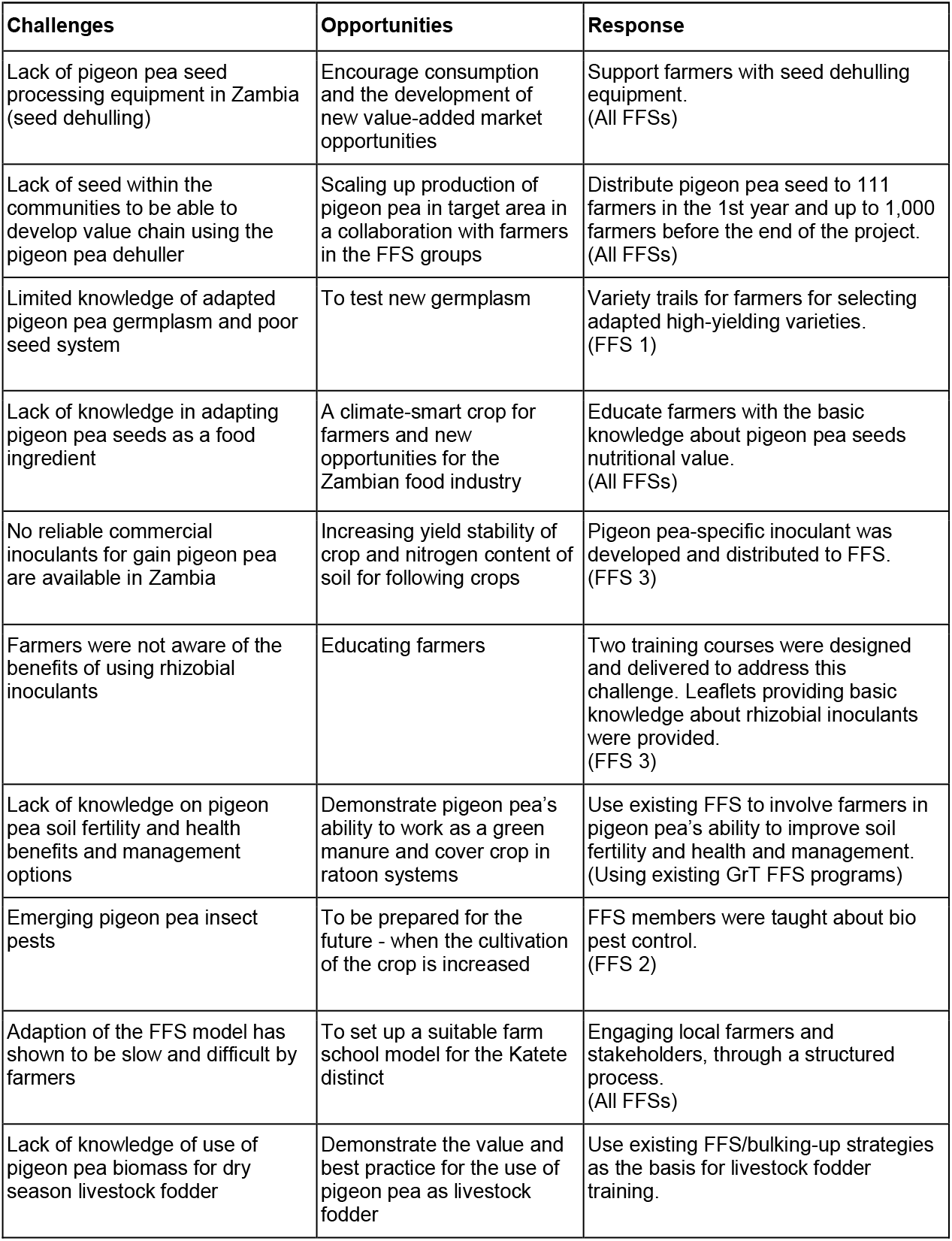
Challenges and opportunities of pigeon pea in the Katete district and Farmer Field School (FFS) responses.

The Katete district has a population of approximately 214,000 people, with the majority (87%) residing in rural areas (CSO, 2022). The Katete district has approximately 43,000 farming families, and about 95% of people in the district rely on agriculture as their main source of income (Hamazakaza et al., 2022). The main crops are maize (*Zea mays*), groundnut (*Arachis hypogaea*), soybean (*Glycine max*), sunflower (*Helianthus annuus*), cowpea (*Vigna unguiculata*), mango (*Mangifera indica*), cotton (*Gossypium hirsutum*) and small gardens, which produce mixed vegetables and fruits. Agropastoral systems dominate the landscape and are rooted in long-standing cultural traditions. Small livestock also play an important role in people’s livelihoods.

As pigeon pea is a relatively new crop in the Katete region, the FFS set out to educate farmers on the characteristics of three commonly available pigeon pea varieties with a view to facilitating participatory variety selection by field school members (**Figure 2a**). The first FFS in the variety trials consisted of two early-medium varieties and one late medium maturity varieties, namely, Muthawajuni (ICEAP 01551), ZPP14 (ICEAP 01514/15), and MPPV2 (ICEAP 00554), respectively.

**Figure 2.**
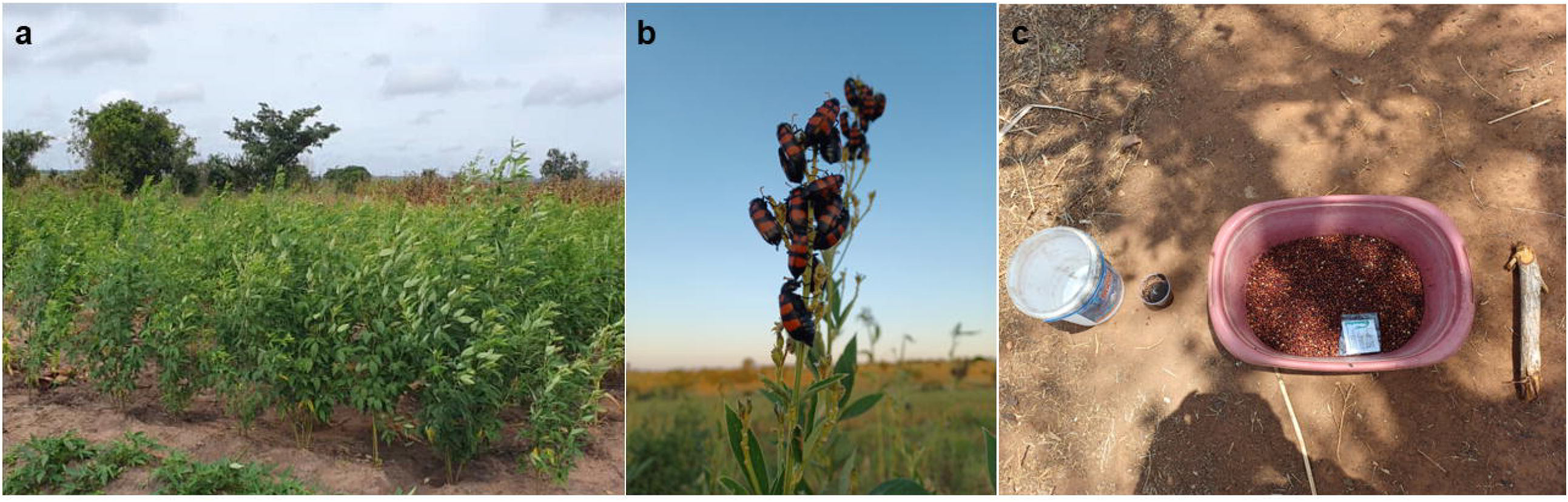
(**a**) Pigeon pea variety trials, FFS1. (**b**) Blister beetles feeding on pigeon pea flowers. (**c**) Training session for farmers on the application of rhizobial inoculation of pigeon pea seeds under FFS 3.

The development of sustainable solutions for pest control is crucial for establishing pigeon pea as a staple crop in the region. The second FFS was structured around low-cost botanical insecticide/repellants for the management of blister beetles (*Mylabris pustulata*), commonly known as the flower or CMR beetle (**Figure 2b**), which is a major pest of legumes, as it eats flowers and young pods. Two locally grown plant species, *Tephrosia vogelii* and *Solanum incanum*, were chosen as the botanical insecticides.

There was a general lack of awareness among FFS farmers regarding the benefits of using pigeon pea-specific rhizobial inoculants (NGO Grassroots Trust and farmers interview). No commercial inoculants are available for pigeon pea in Zambia. To fill these gaps, we delivered experimental scale pea-specific rhizobium inoculants to Zambia for FFS. For this purpose, *Bradyrhizobium japonicum*, a bacterial strain CB756 (HAMBI 1341, University of Helsinki, Finland), was used. The inoculants were produced in Finland (Elomestari Oy, Tornio), and nodulation was tested in a pot experiment at the greenhouse at the University of Helsinki, Finland. After official import permits were obtained from the Zambian Ministry of Agriculture, the inoculants in peat carriers were transferred to Zambia.

The benefits gained from the cultivation of legumes are primarily due to their ability to fix atmospheric nitrogen for use by the crop itself, which leads the crop to produce protein-rich seeds. They also greatly contribute to the soil nitrogen pool, which can benefit neighboring and following crops. The global demand for chemical nitrogen fertiliser and the need for sustainable protein sources have made legumes attractive as part of cropping systems. Appropriate rhizobial inoculants also improve the symbiotic nitrogen fixation and productivity of legumes, particularly under abiotic conditions (e.g., Aserse et al., 2020; Nyaga Njeru, 2020). Therefore, the third FFS focused on assessing the potential of rhizobium inoculants specific to pigeon pea, which is not well documented in the region but has vast potential, as indicated by studies performed in India (Ragendran et al., 2008).

A major challenge is the risk for poor handling, storage, or transportation of inoculants by farmers, which could reduce their viability and effectiveness. To address this concern, FFS were trained (**Figure 2c**) with the following learning objectives: *a*) introduction to rhizobial inoculation; *b*) hands-on training in the effective use of inoculants in the field; *c*) understanding the correct methods for applying inoculants to seeds; and *d*) hands-on training in the proper handling and storage of inoculants.

In 2023, we had 25, 31, and 20 participants in Chimtende (10 F, 15 M), Makwenda (10 F, 21 M), and Greyer (11 F, 9 M), respectively. In 2024, the second batch of rhizobial inoculants was delivered to lead farmers at all three FFS sites, and training was conducted again. The inoculants were handed to 111 farmers across FFS.

Communities were asked if they had other issues that needed to be added to the curriculum for pigeon pea, but because this is a new crop, communities were happy to learn on these three key points. To allow for the potential of generating high-quality scientific data, with the goal of sharing learning outcomes with stakeholders and informing future project directions, these field schools were designed with three replications and a control to allow for meaningful statistical analysis.

During meetings, farmers were presented with the logic surrounding each field school, and the community was asked to select the members whom they thought would be able to manage the field schools to a satisfactory degree. If those people agreed, they were then trained in exactly what needed to be done, given field guides, seeds and any other inputs they needed, such as tape measures, inoculants, etc. Farmers were given the contact details of the trainer and encouraged to communicate any challenges for quick resolution.

The evaluation of the field schools was performed by encouraging the FFS group members to help the host farmers plant if they had time and visit the field school during the season. The NGO Grassroots Trust, which works with these farmer groups, uses an FFS model where farmers are not given regimented assignments, as is commonly done in farmer field schools, which require the group to meet every week. This is a response to well-documented feedback on FFS over the years, where farmers report that the traditional field school approach took too much time from their already busy work schedule, and this presented a significant opportunity cost (Manoj and Vijayaragavan, 2015; Waddinton et al., 2014). The decision to meet is made by the groups themselves. Like in current field schools in Zambia, the facilitator of the FFS is usually present at four specific times during the season, namely, preseason for training and handing out FFS inputs, once during the mid-vegetative stage of the crop, once during the reproductive cycle and finally for after harvest and gross margin analysis. During these four visits, reflections are performed with the group according to observations and results. During the visits, the facilitator inspected the demonstration fields, recorded information on the crop status and yield, and noted any issues reported by the farmers. These visits involved informal interviews, where the facilitator observed basic agronomic conditions and cultivation challenges, engaged with farmers based on these observations, and offered relevant advice. All observations were documented and qualitatively analysed from both advisory and agronomic perspectives. Additionally, unlike other FFS approaches, this program focuses on multi-year crop cycles rather than single cycles, as these take into account differences in climate and markets across seasons. This approach is also important with respect to observing long-term changes in soil fertility and crop performance, a key area of intervention for this NGO. Decisions to meet outside these four times are made by the group members to resolve internal issues or make observations at specific times, e.g., drought occurrence or pest outbreak, if these events do not coincide with the facilitator being present. The reduced cost of this approach over other FFS methodologies offers the potential to reduce barriers to entry into the implementation of FFS over their traditional program structures, as these barriers are well reported to be limiting owing to their associated high cost (Waddinton et al., 2014).

As part of the overall approach of educating farmers on pigeon pea management, farmers were given small amounts of seed for on-farm exposure to this new crop and as a means of ‘bulking up’ pigeon pea seed grain to begin developing a local value chain around pigeon pea processing/dehulling. Over the first two cropping seasons, i.e., the 2023/24 and 2024/25 rainy seasons, 76 (45 M, 31 F) and 111 (50 M, 61 F) farmers, respectively, received pigeon pea seeds under this part of the program. In 2024, the plan was to cultivate approximately 12 hectares of pigeon pea using the seeds distributed to the farmers.

## FINDINGS

The participation of farmer groups has several dynamics, depending on each community. Examples brought up during FFS meetings and conversations with group members revealed that the lack of guaranteed market and pricing (compared with conventional main crops such as maize), two consecutive failed agricultural seasons, and the discovery and development of a gold market in Makwenda and Chimutende seem to have led farmers to be hesitant to allocate time and resources to growing this new crop. In Greya, participation has improved, which could be attributed to Tikondane Community Centre (TIKO) having a long-standing relationship with local farmer groups. TIKO is a community centre run by and for the farmers in Katete (https://www.tikondane.org/). A contributing factor here was that TIKO agreed to host the seed processing equipment, a message that encouraged the farmers to take part in the FFS program.

### Erratic rainfall distribution and extreme weather conditions

During the 2023/24 farming season, Zambia faced a severe drought emergency that resulted in widespread crop failure **(UN Zambia, 2024)**, including the collapse of our pigeon pea FFS and bulking in the Katete district. Based on the visits made to FFS and information obtained from the farmers, most of the 76 farmers failed to establish pigeon pea successfully, and those who did, lost the crop to severe drought at the end of the season.

Second-year erratic rainfall at the beginning of the season also led to the lateness of planting for field schools and for scaling fields. This phenomenon was exemplified by the fact that these farming communities had two years of failed crops (drought 2023/24 and soybean rust disease 2022/23), so they were, understandably, focusing on planting crops they knew that had a secure market for, i.e., maize, soybean, sunflower, and groundnut. Farmers described how they had planted some of their main fields three times due to poor germination at the beginning of the season, and this issue was not limited to pigeon pea. This is particularly relevant to pigeon pea, as early planting is essential to reduce the risk of damage from free-ranging livestock.

In the 2024/25 season, a total of 111 farmers in Katete District received pigeon pea seeds as part of the learning/bulking-up seed distribution initiative. Among the three participating sites, overall, 24 of the 111 farmers (approximately 22%) established pigeon pea crops (**Figure S1**).

### Lack of a value chain

Most farmers doubted that there was a market for pigeon peas in the region. Zambia has found a ready market for pigeon peas in India and wants farmers to capitalise on the opportunity. To date; despite having a small (approximately 100 tonnes/year) local market for processed pigeon peas, Zambia does not have any processing equipment for pigeon peas. As a result, all the processed pigeon pea dahl consumed in Zambia is imported. To support the development of the pigeon pea value chain, the Zambia for Agroforestry, Biodiversity and Climate (Z4ABC) project facilitated the importation of a small-scale dehulling machine from India to Zambia in early 2025. As a result, farmers in Katete now have access to a dehuller. This was perceived as an important improvement in the value chain.

### Lack of developed seed systems for pigeon peas

The program awaits germplasm from ICRISAT to begin making initial crosses and launching a formal breeding program. The Catholic Relief Services (CRS) has been actively involved in the distribution of pigeon pea seeds to farmers over the last decade, sourcing seeds from ZamSeed and ICRISAT. Most of the seeds originated from Malawi. Most of those varieties are mid-to-late maturing varieties, e.g., MPPV2. The later-maturing varieties tend to be eaten by free-ranging livestock in the field after the month of June. Several pigeon pea germplasm have been developed for Eastern and Southern Africa, many of which are potentially adaptable to Zambian agro-ecological conditions (Kaoneka et al., 2016). The potential of selecting suitable lines to improve the dehulling efficiency and market acceptability and pricing is also of interest.

Currently, the majority of the pigeon pea germplasms available in Zambia are vegetable types, which typically have relatively large seeds, relatively low dehulling efficiency and relatively low market prices in India.

### Pigeon pea at risk from free-grazing livestock

Farmers were concerned about the survival of pigeon pea in their fields from May onwards, as this is when livestock owners stop herding their free-range goats and cattle. Since pigeon pea matures around this time, the crop becomes highly vulnerable to grazing. As a result, farmers are only able to harvest once, even though the crop has the potential for up to three harvests per year. This also means that they must replant pigeon pea every season. Fencing their fields is not a viable solution because of labor, financial and traditional land ownership structure constraints. To mitigate the risk of livestock damage, some farmers plan to plant pigeon pea in their home gardens, where they can better protect it from destruction.

### Strengthening farmer engagement through communication

Communication and support to farmers was found to be critical for the success of FFS. Communication and support for FFS could be improved, as groups demonstrated poor communication with the consultant/trainer regarding on-the-ground issues related to the field schools and the scaling up of pigeon pea production. The consultant made several efforts to be given a precise picture of the situation on the ground but only came to realise that things had not gone well during the midseason visits for both years. The exception to this was in Greya in 2025, where communication from one field school host was good and the field school was established successfully and to a good standard. Therefore, besides communication, it is important to make observations on the field to establish a good overview of the situation at each moment. FFS design, field support, and monitoring and evaluation are the main quality indicators that need to be followed by FFS (van den Berg et al., 2021b). FFS are not alternatives to existing systems. However, certain principles of FFS could be selected and incorporated into various systems, such as agricultural extensions (Anandajayasekeram et al., 2007). In our FFS experience, monitoring and evaluation by local or government extension staff could have improved the success of the FFS.

## DISCUSSION

Using the described FFS methodology, where emphasis is placed on the farmer group to carry out the steps toward establishing and maintaining the field school sites and because so much of the learning is dependent on the field itself, there must be buy-in from the FFS group. In this particular case, where we were preempting the production of a new crop, farmers need to have confidence in the end market for the product, pigeon pea dahl.

This was illustrated by the Greya community, in terms of numbers (66 farmers receiving bulking-up seed compared with 24 and 21 for the other communities) who, despite knowing that the dehuller had not arrived, were more closely linked to the potential market through affiliation with TIKO and therefore the prospect of the end market.

The dynamics surrounding farmers’ buy-in through interest and participation in FFS are diverse and varied (Bakker et al., 2020; van den Berg et al., 2020). Taking these considerations into account from the onset of the program will lead to poor participation and implementation. This approach requires efficient and timely communication between facilitators and FFS groups to be effective. Inadequate communication led to suboptimal performance in all but one of the FFS sites in the second year. Therefore, our case study suggests that thorough communication and observing the FFS is critical to ensure that the FFS will reach desired outcomes.

Collaboration with partner organisations could help with the effective establishment of FFS sites, but only if these organisations have the capacity to oversee field operations to the standard needed. The drawback of relying on partner organisations for support is that the FFS hosts invariably wait for extension officers to arrive rather than planting the crop themselves. Late plant establishment is particularly important for pigeon pea, as it needs to be planted early in the season so that the maturity of the crop does not coincide with that of free-ranging livestock.

FFs and their demonstration plots can be a powerful tool to increase rural livelihoods and improve food and nutrition security especially when introducing and upscaling a new crop in a region. However, to valorise the field schools, it is critical to ensure that they operate efficiently.

## ACKNOWLEDGEMENTS

We thank our Zambia CIFOR colleagues, Maimbo Malesu, Chilala Ndeke, Wellington Chazya, Dr. Maarit Kallio, Dr. Nicholas Hogarth, Chipo Chisonga, and Dr. Patricia Masikati, for their kind support during this activity. We also thank all the farmers and communities, including TIKO (Tikondane Community Centre), who participated in the FFS activities. Special thanks to Elke Kroeger-Radcliffe (TIKO) for her kind support for FFS. Many thanks to the Ministry of Agriculture, Kabanghe Masenga (District Agriculture Coordinator), Ziko Kahenge (Senior Agriculture Officer), Theresa Bwalya (Block Officer), Moses Banda (Agricultural Assistant), Peter Bwalya (Agricultural Assistant) and Zambia Agricultural Research Institute (ZARI) – Msekera Research Station in Chipata, and the legume breeder Lutangu Makweti for his valuable suggestions and for providing pigeon pea seeds. Petri Leinonen (Elomestari Oy) is acknowledged for helping with the rhizobial inoculants.

## AUTHOR CONTRIBUTIONS

H.K. and S.C. wrote the original draft. All the authors wrote, reviewed and edited the manuscript. All the authors read and agreed with the published version of the manuscript.

## ETHICS APPROVAL AND CONSENT TO PARTICIPATE

Ethics approval was not required for the present study. All farmers participated voluntarily to the FFSs, and they had to indicate their consent to participate in the pigeon pea activities. Before joining the FFSs, the farmers were explained the purpose of the pilot, and they had the possibility to ask questions before deciding whether to participate in the pilot. The farmers had an option to discontinue in the pilot at any time without having to explain the reason.

## DATA AVAILABILITY

The datasets developed during the current study are available from the corresponding author upon request.

---

**Figure S1.**
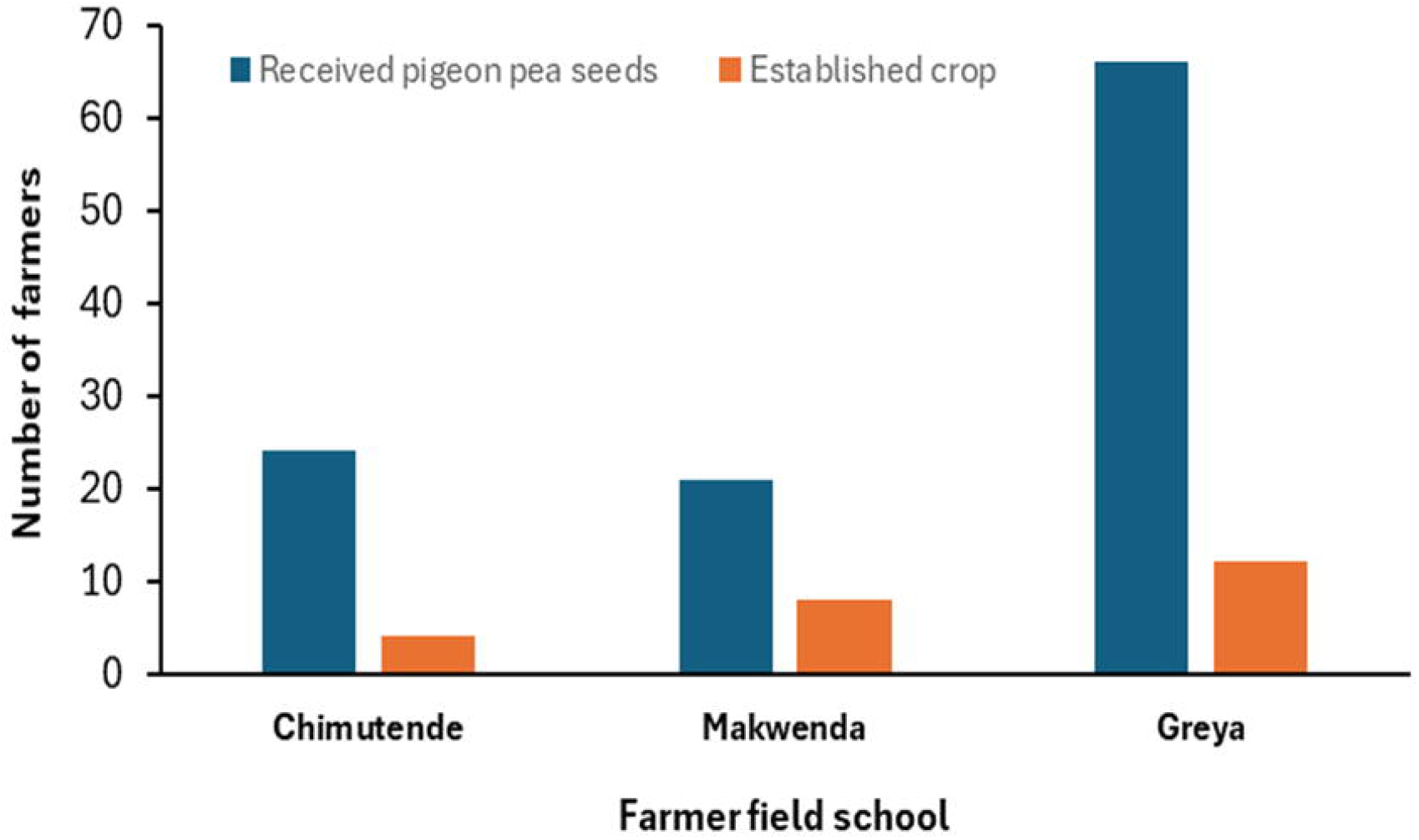
Pigeon pea learning/bulking-up seed distribution and number of farmers who planted in the Katete FFSs 2024/25 season.

